# Particle-associated and free-living bacterial communities in an oligotrophic sea are affected by different environmental and anthropogenic factors

**DOI:** 10.1101/2020.04.20.051391

**Authors:** Dalit Roth Rosenberg, Markus Haber, Joshua Goldford, Maya Lalzar, Dikla Aharoonovich, Ashraf Al-Ashhab, Yoav Lehahn, Michael D. Krom, Daniel Segrè, Laura Steindler, Daniel Sher

**Affiliations:** Department of Marine Biology, Leon H. Charney School of Marine Sciences, University of Haifa, Israel; Bioinformatics Program, Boston University, USA; Bioinformatics Support Unit, University of Haifa, Israel; Microbial Metagenomics Division, Dead Sea and Arava Science Center, Masada, Israel; Department of Maritime Geosciences, Leon H. Charney School of Marine Sciences, University of Haifa, Israel; Morris Kahn Marine Research Station, Department of Marine Biology, Leon H. Charney School of Marine Science, University of Haifa, Haifa, 3498838, Israel

**Keywords:** Eastern Mediterranean, 16S amplicon sequencing, Particle-associated, Free-living, Seasonality, anthropogenic pollution, phytoplankton

## Abstract

In the oceans and seas, environmental conditions change over multiple temporal and spatial scales. Here, we ask what factors affect the bacterial community structure across time, depth and size fraction during six seasonal cruises (two years) in the ultra-oligotrophic Eastern Mediterranean Sea. The bacterial community varied most between size fractions (free-living vs particle-associated), followed by depth and finally season. The free-living (FL) community was taxonomically richer and more stable than the particle-associated (PA) one, which was characterized by recurrent “blooms” of heterotrophic bacteria such as *Alteromonas* and *Ralstonia*. The heterotrophic FL and PA communities were also correlated with different environmental parameters: depth and phytoplankton correlated with the FL population, whereas PA bacteria were correlated primarily with season. A significant part of the variability in community structure could not, however, be explained by the measured environmental parameters. The metabolic potential of the PA community, predicted from 16S amplicon data, was enriched in pathways associated with the degradation and utilization of biological macromolecules, as well as plastics, other petroleum products and herbicides. The FL community was enriched in pathways for the metabolism of inositol phosphate, a potential phosphorus source, and of polycyclic aromatic hydrocarbons.

**Originality – Significance Statement:** Marine microbial populations are complex and dynamic, and the environmental drivers of the structure and function of these communities are mostly unclear. Specifically, marine microbial communities change over time, over depth and between particle-associated and free-living size fractions, yet the relative importance of each of these axes of variability is unclear. Our results highlight fundamentally different population dynamics between free-living and particle-associated marine bacteria: free living populations were more similar between seasons, whereas particle-associated populations were highly variable and exhibited “blooms” of specific clades of heterotrophic bacteria. We also suggest that the environmental conditions often measured as part of oceanographic cruises are not enough to explain most of the variability in microbial population structure. We speculate that organismal interactions and the presence of anthropogenic pollution may be also be important yet under-sampled drivers of oligotrophic marine microbial communities.

## Introduction

To a microorganism, the marine environment presents a rich and ever-changing tapestry of conditions and potential niches. On scales of microns to millimeters, which can be traversed by motile bacteria within minutes, various types of particles and gels provide carbon- and nutrient-rich hotspots which differ from the bulk surrounding seawater in their chemistry and physics (Azam and Malfatti 2007). Other environmental parameters such as light intensity, temperature and the concentrations of dissolved inorganic and organic nutrients change markedly with depth over scales of tens to hundreds of meters, particularly when the water column is stratified (Karl 2007). Depending on the structure of the water column and the vertical movement of macro-organisms and sinking particles, bacteria can traverse such distances (both up and down) on scales of hours to days (e.g. (Grossart et al. 2010, Mestre et al. 2018)). Finally, changes in environmental conditions can occur over spatial scales of hundreds of kilometers or temporal scales of weeks to months. For example, seasonal changes in weather or geographic changes in environmental conditions are driven by global climate patterns (Giovannoni and Vergin 2012). Seasons affect the physical structure of the water column and, through mixing of water masses, control the injection of inorganic nutrients into the photic zone. Indeed, microbial populations in the oceans differ over multiple spatial and temporal scales (Fuhrman 2009, Martin-Platero et al. 2018, Salazar et al. 2019). While our understanding of the processes that shape marine microbial communities over these scales has been steadily increasing (Karl et al. 2002), many questions remain unanswered. For example, changes in microbial population structure have been documented between different size fractions (e.g. (Mestre et al. 2017)), along a depth gradient (DeLong et al. 2006) and over seasonal cycles (Ward et al. 2017), yet few studies have addressed these spatio-temporal axes of variability together. Which of these environmental factors has a more pronounced effect on microbial populations? Are changes in heterotrophic microbial populations driven primarily by variation in “a-biotic” conditions, or do other factors affect population structure, for example biotic interactions with co-occurring phytoplankton? Can we predict which genetic traits or metabolic functions underlie changes in community structure along these gradients of variability?

To begin addressing some of these questions, we characterized the environmental conditions, the phytoplankton community, and the bacterial community structure across multiple spatio-temporal scales in the Levantine Basin of the Eastern Mediterranean Sea (EMS). Despite being an inland sea, surrounded by ^~^480 million people (Bleu 2009), the open waters of the Mediterranean, and particularly the EMS, are oligotrophic to ultra-oligotrophic (Fig. 1A, (Berman et al. 1985)). Nutrient concentrations in the photic zone are typically very low (close to the level of detection), as are chlorophyll and particulate carbon, and the photic zone may extend quite deep, with a dominant Deep Chlorophyll Maximum (DCM) often observed as deep as ^~^140m (Berman et al. 1985, Krom et al. 2005). Phytoplankton in the EMS are often phosphorus-limited or nitrogen and phosphorus co-limited (Krom et al. 2005, Thingstad et al. 2005). Pico-cyanobacteria such as *Prochlorococcus* and *Synechococcus* are the numerically dominant phytoplankton, although photosynthetic pico-eukaryotes are also common and potentially dominant in terms of production or biomass (Man-Aharonovich et al. 2010). The deep waters of the EMS are much warmer than those of many oceanic regions, with the temperature never dropping below ^~^12°C. The deep waters are mixed into Levantine Intermediate waters and these combined waters exit the Eastern Mediterranean (EMS) at the Straits of Sicily, eventually exiting the Mediterranean through the straits of Gibraltar and affecting large parts of the Atlantic Ocean. Recent studies have shown that several water masses of the EMS are warming and becoming more saline at a rate significantly higher than the average global trend predicted by the IPCC (Ozer et al. 2017). Thus, while the EMS exhibits conditions reminiscent of those in the major oceanic gyres (Powley et al. 2017), it is a potentially useful “natural laboratory” to understand the effect of environmental conditions, including climate change, on microbial processes.

**Fig. 1:**
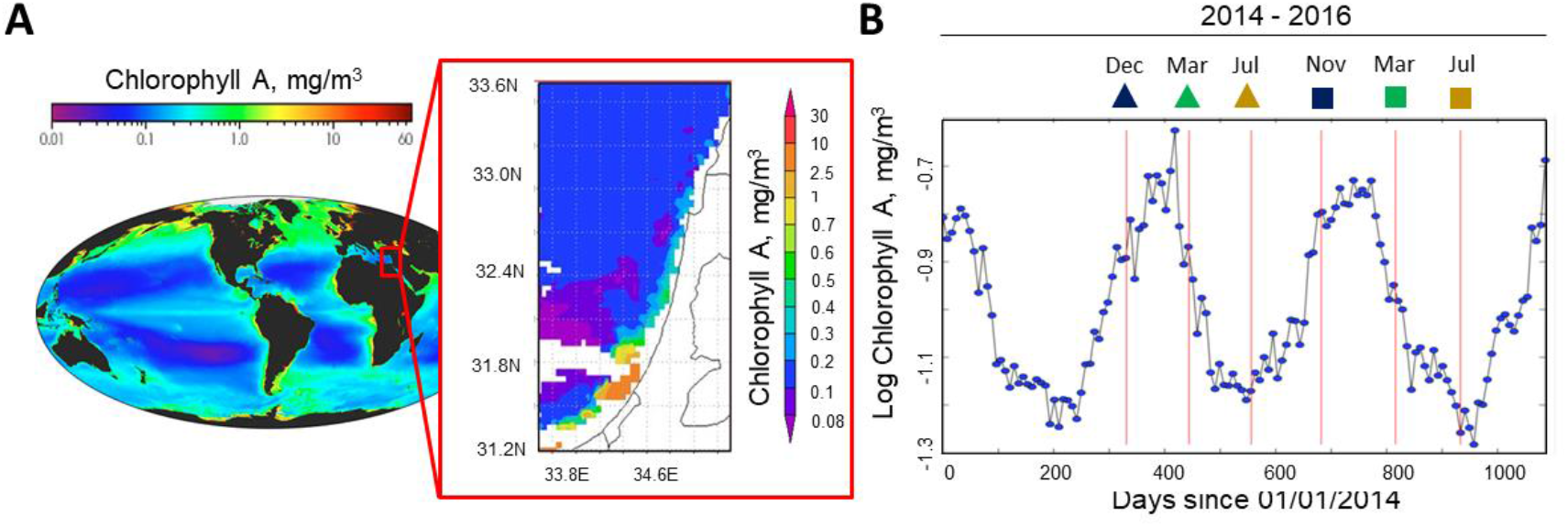
Seasonal patterns in the oligotrophic Eastern Mediterranean Sea (EMS). (A) The EMS is ultra-oligotrophic, with conditions comparable to major ocean gyres. Maps are shown of the surface chlorophyll *a* concentrations (in mg/m^3^) between 1997-2004 (world image, downloaded from the NASA website) and along the Israeli coast (average March 22nd-29th, 2015, from the MODIS-aqua 4km data set). (B) Annual dynamics of sea surface chlorophyll from satellite data, showing the timing of the six research cruises during fall (blue), spring (green) and summer (red).

To obtain a dynamic view of how environmental parameters at multiple scales affect the microbial population structure, we present and interpret physical, chemical and biological information from six cruises to a pelagic station in the EMS. Station n-1200 is ^~^25 nautical miles from shore, where the water column depth is 1200m. The cruises were performed during three seasons (fall, spring and summer) of two consecutive years. Samples were collected from five discrete depths (surface, ½ DCM, DCM, 200m and 500m) and separated into different size fractions, >11μm and 5-0.22μm, representing particle-associated and free-living microbes, respectively. These data allow us to describe and interpret the natural changes in microbial populations in the EMS over different scales: (i) particle associated *vs*. free-living, (ii) over depth, and (iii) over seasons.

## Results and discussion

### Physical, chemical and biological properties of the EMS during the cruises

During 2014-2016, a clear seasonal cycle in sea surface Chlorophyll was observed in the EMS, with maxima in winter and early spring and minima in summer, consistent with previous *in-situ* measurements (Fig. 1B, (Azov 1986, Raveh et al. 2015)). Our sampling cruises were timed to coincide with the increase in surface chlorophyll during fall, the subsequent decrease phase during spring, and the stable, ultra-oligotrophic summer conditions. The changes in surface chlorophyll were associated with changes in the structure of the water column, which was highly stratified during summer, with the depth of mixing increasing during fall and peaking during early spring (Supporting Information Fig. S1). Macronutrient profiles show that the surface waters, down to ^~^200m, were poor in nitrate+nitrite and phosphorus with the lowest values in July compared to fall and spring (Supporting Information Fig. S2). A deep chlorophyll maximum (DCM) was observed in all cruises apart from the November 2015 one, and was generally shallower during fall and spring (70-90m) and deeper during the summer (90-140m, Supporting Information Fig. S1).

While clear seasonal patterns were observed in the surface chlorophyll, more complicated patterns were observed in the composition and relative abundance of different phytoplankton groups within the photic zone. Based on flow cytometry counts, during most of the year, *Prochlorococcus* were the numerically most abundant phytoplankton, particularly deeper in the water column (½ DCM and DCM, Supporting Information Fig. S3). *Synechococcus* were usually more abundant at the surface (10m and occasionally ½ DCM). Consistent with the flow cytometry counts, divinyl-chlorophyll *a* (a pigment which is unique to *Prochlorococcus*) peaked at the DCM, yet never comprised more than ^~^12.5% of the total chlorophyll concentration, suggesting that other phytoplankton may be more dominant in terms of biomass (Supporting Information Fig. S4). Indeed, 19’-hex-fucoxanthin (“19-hex”), a pigment characteristic of prymnesiophytes (Jeffrey et al. 2005), was relatively abundant at the DCM, particularly during the first year. While 19’-but – fucoxanthin (“but”) and fucoxanthin (“fuco”) were also observed, we cannot conclude that diatoms were present at high abundances, as these pigments are also found in prymnesiophytes. Peridinin, which is characteristic of dinoflagellates, was observed mainly in the fall of both years and the spring of the second year, at the ½ DCM and DCM depths. These results are consistent with previous studies suggesting that pico-eukaryotes, and particularly coccolithophores (which are prymnesiophytes), are an important part of the phytoplankton community, which is dynamic across time and depth (Man-Aharonovich et al. 2010). These results also suggest that non-seasonal (potentially aperiodic) dynamics may influence phytoplankton dynamics (Karl et al. 2002).

### Overview of the changes in bacterial population structure across size fraction, depth and season

As shown in Figure 2 and Supporting Information Fig. S5, clear differences were observed between the particle-associated bacterial communities (PA, >11μm) and the free-living ones (FL, 5-0.22μm). Several samples representing the intermediate size fraction (11-5μm) were also analyzed, and these revealed a population structure somewhat similar to both PA and FL fractions, in agreement with previous studies suggesting a continuous shift in population structure across different size fractions ((Mestre et al. 2017, Mestre et al. 2017), Figure 2A, B). The FL and PA populations both varied with depth, although the changes with depth in the PA community were less pronounced (see details below). No clear clustering of the populations by season was observed in the NMDS ordination (Figure 2B). Indeed, the Bray Curtis dissimilarity was highest between FL and PA communities, followed by the difference with depth, with the lowest variability observed between seasons (Figure 2C). Thus, in the open EMS, size fraction and depth are likely the most important drivers of microbial community composition.

**Figure 2:**
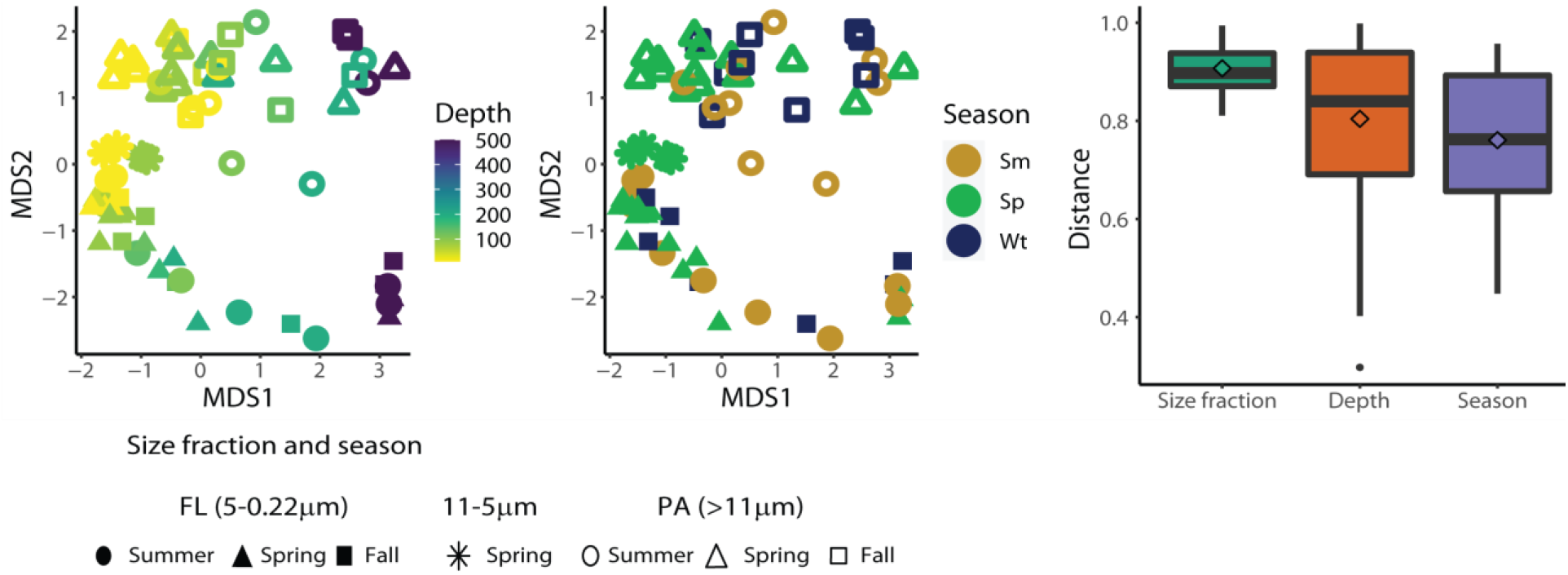
Effects of size fraction, depth and season on bacterial population structure. (A, B) Nonparametric Multi-Dimensional Scaling (NMDS) plot of the bacterial populations, colored by depth (A) and by season (B). Full shapes are FL, open shapes are PA, with the two groups clearly separated along NMDS axis 2. Both FL and PA communities change over depth along NMDS axis 2, but do not cluster by season. NMDS stress = 0.11. (C) Bray-Curtis dissimilarity is highest with size fraction, followed by depth and season. The dissimilarity was calculated within groups, e.g. comparisons of depth were performed within the same cruise and size fraction. The means of the size fractions are all statistically different (t-test, p<0.001, (Fagerland 2012)).

The differences between FL and PA communities were lower at the surface, increasing with depth (Fig. 2, Supporting Information Fig. S6A). This is consistent with a mechanism whereby many particles originate in the surface layer, either formed autochthanously or added as atmospherically derived particles (e.g. dust (Herut et al. 2002)). The particles are colonized primarily by surface layer bacteria, with the exchange between FL and PA populations becoming less common with depth (Mestre et al. 2018). At a coarse grained phylogenetic resolution, the FL population was dominated primarily by 16S sequences belonging to α-proteobacteria from the SAR11 clade (*Pelagibacter*), which comprised up *to* ^~^40% of the sequences in the FL fractions, mostly in the surface waters, (Supporting Information Fig. S5). Other abundant heterotrophic lineages were SAR86 (γ-proteobacteria, up to 15% in surface waters during summer and spring), SAR 406 (Marinimicrobia, up to 8% in deeper waters) and SAR 202 (Chloroflexi, primarily intermediate depths, up to 12%). Conversely, the PA communities were rich in γ-proteobacteria and Planctomycetes, as well as occasionally β-proteobacteria, δ-proteobacteria or Verrucomicrobia, phyla previously described as associated with marine particles from the Atlantic (Milici et al. 2016), Baltic (Rieck et al. 2015) and Mediterranean Seas (Mestre et al. 2017, Mestre et al. 2017). Notably, while pico-cyanobacteria such as *Prochlorococcus* and *Synechococcus* were primarily observed in the FL fraction, they also comprised as much as ^~^6% of the 16S reads from the PA communities, and up to ^~^21% of the communities in the size range of 11-5μm (Supporting Information Fig. S5, Supplementary excel file). *Prochlorococcus* and *Synechococcus* are primarily free-living cells, however, previous studies have suggested that pico-cyanobacteria do contribute to particulate sinking fluxes, although the magnitude of this process is unclear, and may depend on the specific oceanographic conditions (Richardson and Jackson 2007, Lomas and Moran 2012, De Martini et al. 2018). The presence of *Prochlorococcus* and *Synechococcus* cells in the PA fractions suggests a potential involvement of these clades in sinking fluxes in the EMS.

Within the FL fraction, the population structure clearly changed with depth, with the main differences seen between the communities from the photic zone (here defined as down to 200m) and the mesopelagic (200m and 500m, Fig. 2A, Supporting Information Fig. S5, S6B). The dissimilarity among sampling depths was higher during stratified seasons than during spring, consistent with a partial homogenization of the water column by winter/spring mixing (t-test, p<0.001, Supporting Information Fig. S6B). As expected from other studies, cyanobacteria and Bacetroidetes were relatively more abundant in the photic zone, whereas δ-proteobacteria, Marinimicrobia and Chloroflexi were more associated with deeper samples (Haro-Moreno et al. 2018, Mende et al. 2019). The change of the PA community with sampling depth was weaker, observed in the NMDS analysis (Figure 2) but not in the one-dimensional clustering (Supporting Information Fig. S5), and no clear differences with sampling depth were observed (Supporting Information Fig. S6C). Instead, the PA community seemed to partition into samples dominated by γ-proteobacteria (primarily *Alteromonas*), and those where Planktomycetota, Bacteroidetes or β-Proteobacteria were more dominant (Supplementary Figure 5 and see below).

In contrast to the clear effects of particle association and depth on microbial community structure, no obvious partitioning by season could be observed in the NMDS or clustering (Figure 2B, Supporting Information Fig. S5). Nevertheless, seasonal differences were observed when the same size fraction and depth were compared (Supporting Information Fig. S6B, C, D). The seasonal differences were significantly larger in the PA compared to the FL populations, suggesting that qualitatively different particles may be found at different seasons. Surprisingly, within the FL population, seasonal variability was lower at the surface and ½ DCM compared with the deeper samples. We initially expected seasonal differences to be highest at the surface, reflecting major changes in sea surface temperature, which has been suggested to be major driver of microbial community structure and function (Sunagawa et al. 2015). The higher inter-seasonal differences detected at intermediate depths (100-200m) suggest that factors other than temperature are causing seasonal changes in the populations at these depths. These may include the quality and intensity of light supporting variable deep phytoplankton populations, or the availability of nutrients mixed up from the nutricline (Karl et al. 2002).

### Particle-associated and free-living communities have different strain-level (ESV) dynamics

A detailed inspection of the community structure (Supporting Information Fig. S5, Supporting excel file) suggests that some PA communities are dominated by a limited number of ESVs (exact sequence variants, which represent distinct 16S phylotypes). Indeed, the FL population was more diverse (Shannon and inverse Simpson indices) and more even than the PA ones (Figure 3A). These results are consistent with studies from major ocean gyres and the offshore western Mediterranean (Ghiglione et al. 2007, Mestre et al. 2018), whereas studies from the North Sea, the Baltic Sea and coastal locations in the western Mediterranean suggest that PA communities are more diverse than FL ones (Bižić-Ionescu et al. 2015, Rieck et al. 2015, Mestre et al. 2017). It is possible that the productivity (trophic status) of the community determines whether FL or PA communities are more diverse, perhaps because more productive regions tend to produce larger particles or larger phytoplankton with potentially more micro-niches. The alpha diversity indices of the FL populations, but not the PA ones, increased with depth throughout the photic zone (Supporting Information Fig. S7). This again suggests that, despite the variability in surface temperature, intermediate depths may provide more niches for FL bacteria than the surface waters.

**Figure 3:**
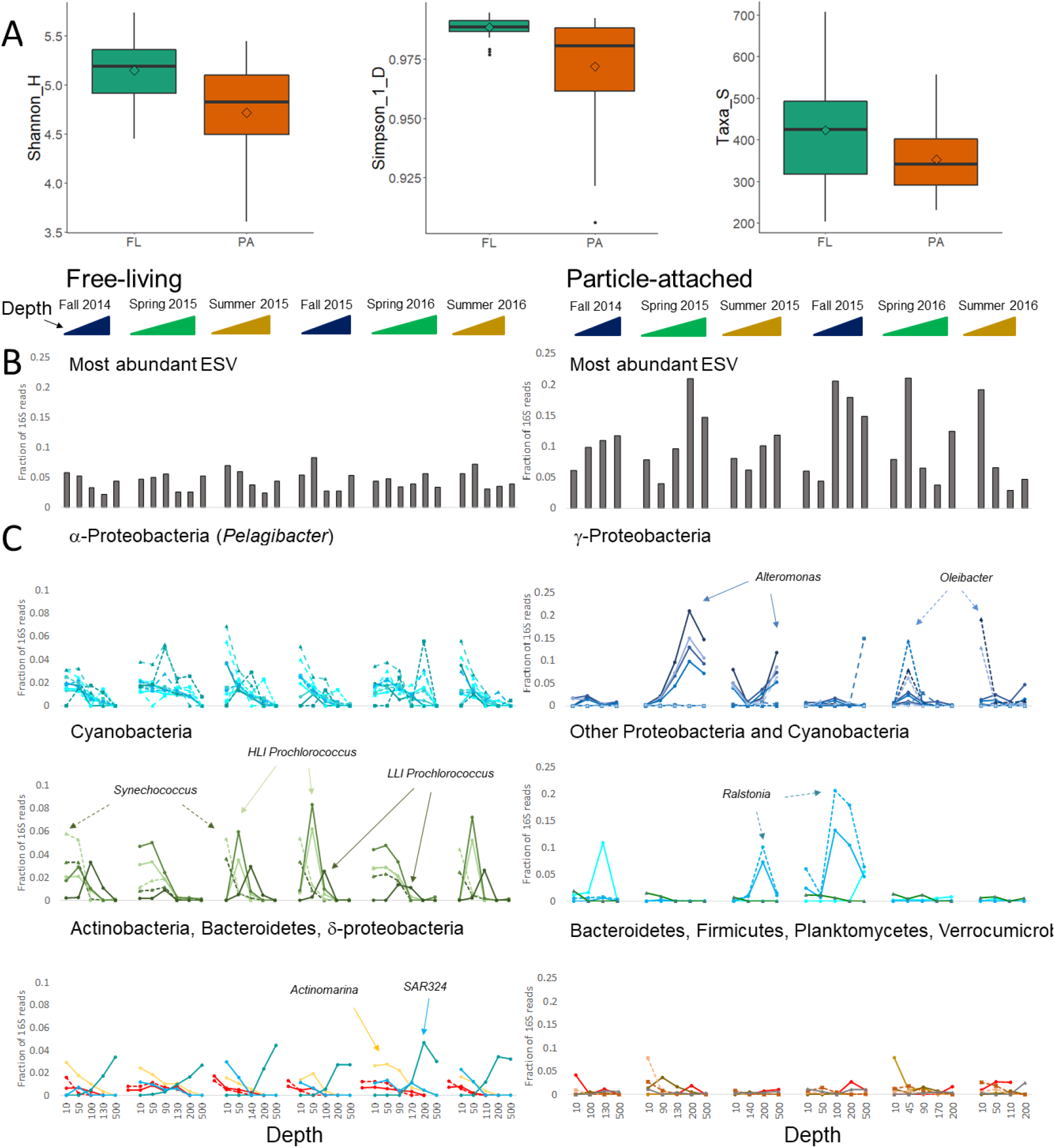
Diversity and ESV dynamics differ between FL and PA communities. In panels B and C, each individual graph shows a depth profile of a separate cruise. (A) FL populations are more diverse than PA ones. Boxes show medians, 25^th^ and 75^th^ quartiles; whiskers show 1.5 times the interquartile range, and outliers are plotted individually. Diamonds show means. All differences are statistically significant (t-test, p<0.001 for comparisons of Inverse Simpson and Shannon indices and p=0.002 for comparison of the number of taxa). (B) Relative abundance of the most common ESV in each sample (cruise, depth and size-fraction). (C) Relative abundance of the 20 most common ESVs across each of the FL and PA populations. For depths with duplicate samples, the mean of the duplicates is shown. ESVs are colored based on their phylogeny using the same color code as in Supporting Information Fig. 5, and specific ESVs mentioned in the text are highlighted by arrows. For detailed information please see Supporting Information Excel file.

Consistent with the higher α-diversity of the FL populations, the most abundant ESVs comprised at most ^~^8% of the 16S reads of the FL population, compared with up to 21% of the reads for in the PA community (Fig. 3B). While these differences could, in principle, be due to more 16S genes per genome in the PA bacteria compared with the FL ones, we also observed many more ESVs (each with at least 100 reads across our dataset) corresponding to the abundant FL clades compared with the PA ones. For example, there were 192, 71 and 60 different ESVs belonging to the abundant FL clades SAR11, SAR86 and SAR406, respectively, compared to 9, 3 and 2 ESVs belonging to the *Alteromonas, Oleibacter* and *Ralstonia* clades that dominated specific PA samples.

The FL and PA communities also differed qualitatively in the temporal dynamics of the most abundant ESVs. The FL communities were relatively similar across cruises, changing primarily with depth. For example, the ten most relatively abundant SAR11 ESVs were all observed in at least one depth from each cruise, albeit with differing temporal and depth patterns (Fig. 3C). Similarly, the five most fractionally abundant cyanobacterial ESVs were also observed in all of the cruises, and their dynamics over time and depth were consistent with the flow-cytometry counts (Supporting Information Fig. S3) and with ecotype-level dynamics observed in other oligotrophic oceans (e.g. (Malmstrom et al. 2010)). In contrast, many of the most fractionally abundant ESVs in the PA populations were highly dominant in samples from one or two cruises, and rare or almost absent in others. For example, four ESVs, all closely related to *Alteromonas macleodii* (a γ-proteobacterium), were very common in the spring and summer of 2015, together comprising up to 65% of the sequencing reads. These same ESVs were found at much lower relative abundances at other sampling times. Similarly, *Ralstonia* (a β-proteobacterium) dominated intermediate depths in summer and fall of 2015, and *Oleibacter* (also a γ-proteobacterium) dominated during the spring and summer of 2016. In many cases, the same ESVs dominated replicate samples from the same time or samples from different depths of the same cruise (Supporting Excel file), suggesting that these “heterotroph blooms” are not limited to individual particles but rather represent a general feature of the PA community at a specific time and depth. A more detailed description of the dynamics of SAR11, *Prochlorococcus, Alteromonas* and *Ralstonia* is presented in the supplementary information.

### Different environmental factors affect the free-living and particle-associated heterotrophic populations

The increase in FL bacterial diversity at intermediate depths, and the “heterotroph blooms” observed in the PA community, prompted us to ask whether the heterotrophic bacterial community structure might be determined, at least in part, by interactions with specific phytoplankton groups. Phytoplankton can affect heterotrophic bacteria by providing particulate niches and sources of organic material, as well as through direct signaling (reviewed by (Amin et al. 2012, Buchan et al. 2014, Cirri and Pohnert 2019, Durham et al. 2019)). To answer this question, we sought for statistical correlations between the heterotrophic bacterial community structure and three matrices of conditions corresponding to those associated with seasonality, depth and phytoplankton community structure, with the latter defined by the ratios of the concentrations of photosynthetic pigments (Supporting Information Figure S4, Supporting excel file). As shown in Fig. 4, the matrix of depth-related parameters explained the largest amount of variability in the FL populations (16%), with water depth and the concentration of NO3+NO2 being significant explanatory variables. The matrices related to season and pigments alone explained little variation, however, the combination of the matrices of pigments and depth had significant explanatory power. Of the four photosynthetic pigments significantly correlating with the FL heterotrophic population structure, two are associated with cyanobacteria, and specifically *Prochlorococcus* (DVChl-a, Chl-b), and one with prymnesiophytes (19’-hex). In contrast to the FL heterotrophic community, season-related factors, and specifically the depth of the mixed layer (MLD), were the only statistically significant determinants of PA heterotrophic communities, with a small but significant interaction with phytoplankton community. Thus, the main factors driving the heterotrophic FL and PA community structures were fundamentally different. In total, only 30-43% of the variation in heterotrophic population structure could be explained by correlations with seasonality, depth and/or phytoplankton community structure, suggesting that other environmental factors, which were not measured in this study, are important drivers of heterotrophic bacterial communities.

**Figure 4:**
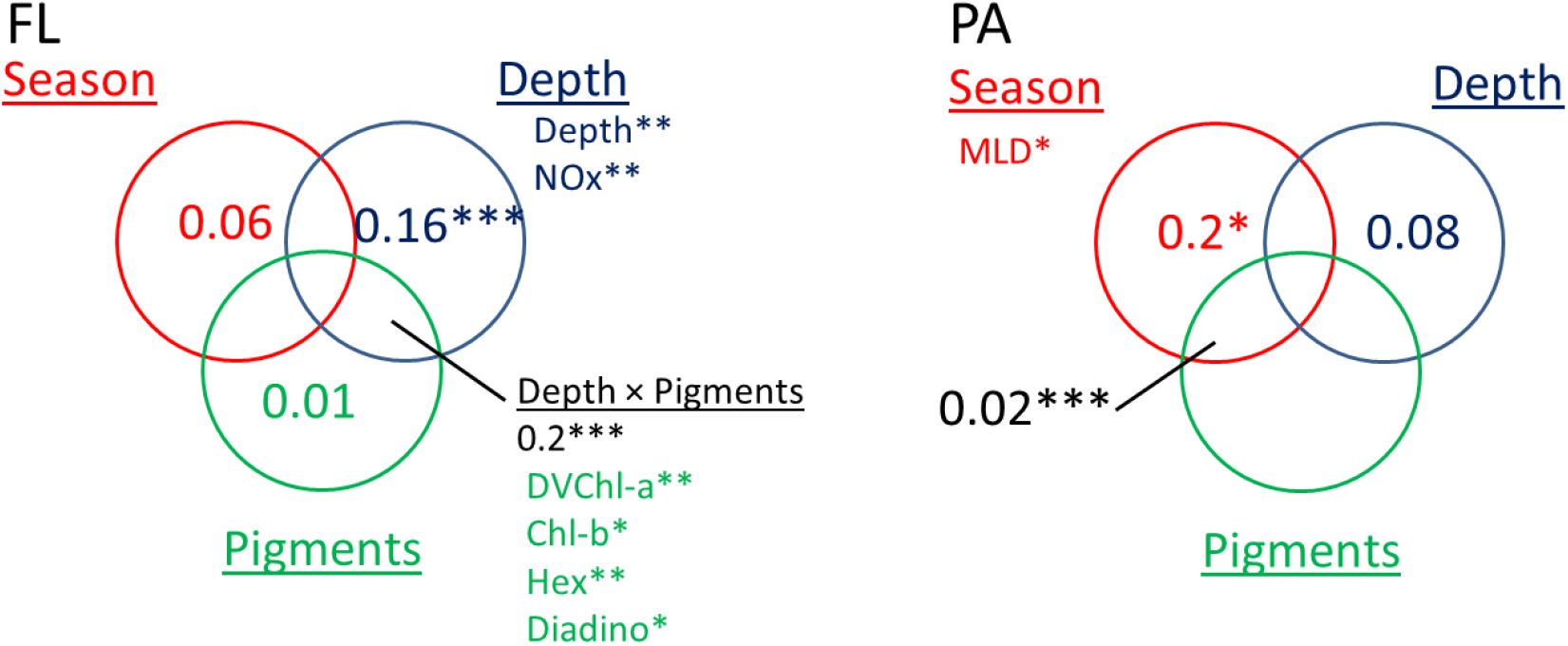
Different environmental factors are correlated with the structure of the PA and FL heterotrophic populations. The results of variation partitioning analysis are shown, with the numbers representing the fraction of the variation explained by each group of environmental parameters. When no numbers are shown, this matrix had no explanatory power (zero or negative). Stars represent statistical significance: * - <0.05, ** - <0.01, *** - <0.001, as estimated using canonical correspondence analysis. When pigments are shown, it is their ratio to chlorophyll a that is significantly correlated with heterotrophic population structure.

### Predicting metabolic functions that might underlie changes between the FL and PA heterotrophic community structure

In this study, we measured multiple environmental parameters known to affect bacterial physiology in the lab, and which could be expected to drive bacterial community structure in the oceans (Supporting Information Figs S1-4). Given that >55% of the variability could not be explained by these factors (Fig. 4), we used the 16S amplicon data itself to raise testable hypotheses as to additional environmental conditions that might drive heterotrophic population structure. We inferred the metagenome of the heterotrophic populations using PICRUSt (Langille et al. 2013), and searched for specific metabolic pathways predicted to be differentially abundant between the heterotrophic PA and FL populations, as the differences between these microbial populations were most clear (Fig. 2, 3). As shown in Fig. 5, many pathways were enriched in the PA compared with the FL heterotrophic populations, consistent with the generally larger genomes and richer metabolic capacities of particle-associated bacteria (Lauro et al. 2009). As expected, many of the pathways enriched in the heterotrophic PA bacteria were related to the degradation of macromolecules that comprise part of phytoplankton cells or zooplankton carapaces (e.g. glycoseaminoglucans, starch, lipids, steroids and amino acids and precursors such as benzoate, Fig. 5). Many other enriched pathways, however, were for the degradation of anthropogenic pollutants. These included pathways for the degradation of organic compounds used extensively in the petroleum or plastic industries, such as caprolactam, styrene, xylene, bisphenol A, ethylbenzene and toluene/nitrotoluene. Other pathways were related to the degradation of herbicides, pesticides or compounds involved in their biosynthesis (e.g. fluorobenzoate, atrazine). Plastics (including micro-plastic particles) are ubiquitous in many marine environments (e.g. (Kwon et al. 2015)). They may potentially be important in the dominantly down-welling EMS, in a similar way to their concentration in mid-oceanic gyres (Powley et al. 2017, Chen et al. 2018). Plastics provide surfaces which bacteria can colonize and potentially utilize (Oberbeckmann and Labrenz 2020). Plastics also leach dissolved organic carbon, which may stimulate bacterial growth (Romera-Castillo et al. 2018), but these leachates can also inhibit the growth of key organisms such as pico-phytoplankton (Echeveste et al. 2010, Tetu et al. 2019). Finally, plastics are also thought to adsorb many other contaminants such as pesticides, leading to a local increase in the latter molecule’s concentration (e.g. (Rios et al. 2010, Chen et al. 2018)). We speculate that the presence of microplastics may select for PA bacteria able to colonize and utilize these carbon sources, while being resistant to toxic leachates and adsorbed contaminants, thus affecting heterotrophic PA population structure.

**Figure 5:**
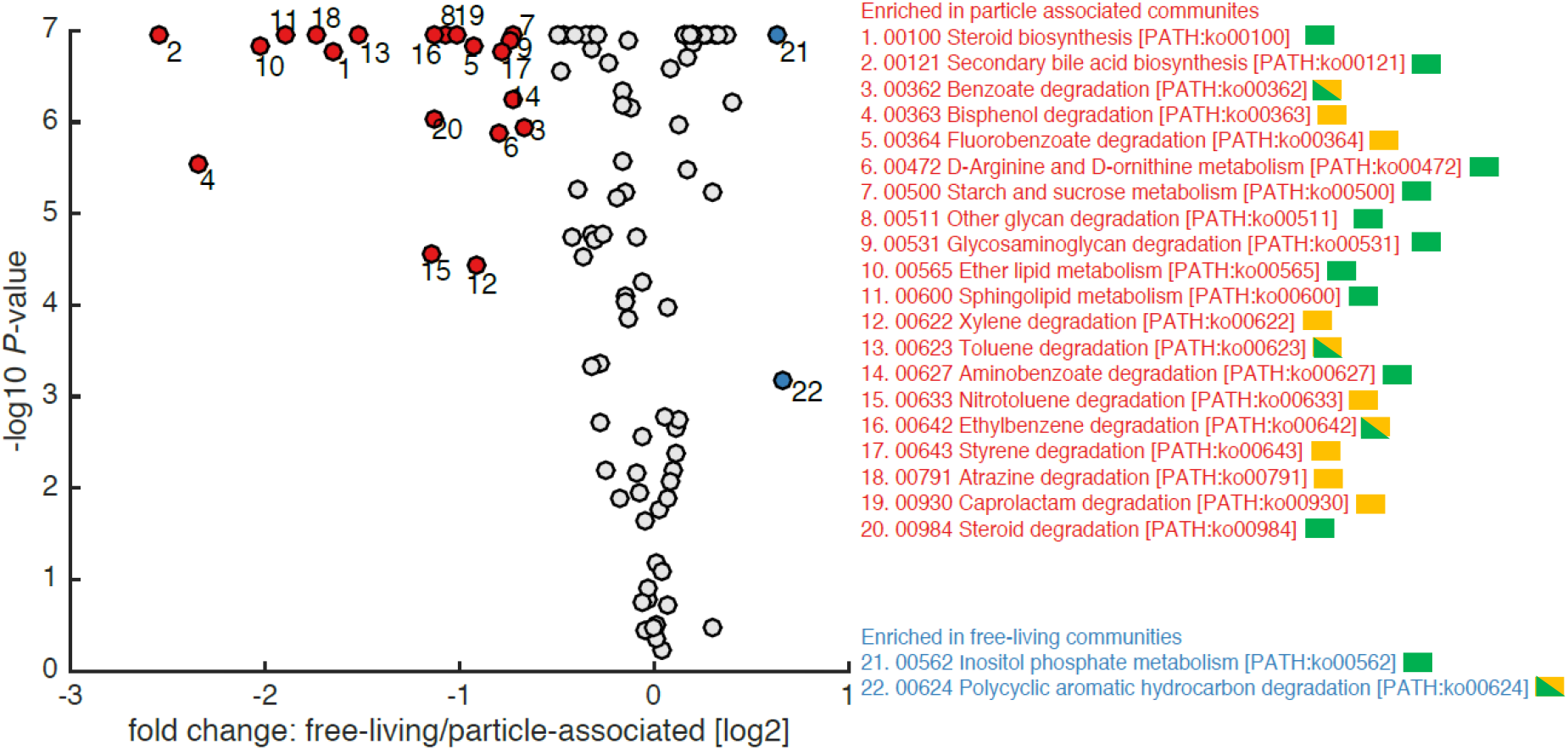
KEGG pathways predicted by PICRUSt to be enriched in the PA (red) and FL (blue) heterotrophic populations. Pathways are ordered by KEGG number and those related to the degradation of animal or plant-derived compounds and of anthropogenic pollutants are marked in green and orange, respectively. Benzoate and ethylbenzene are produced anthropogenically in large amounts but can also occur naturally (in natural petroleum or tar), and thus are marked in both orange and green. Similarly, toluene is used extensively as a thinner and in the petroleum industry, but can also be found in some resins, and is thus marked in both green and orange. The number of pathways enriched in the heterotrophic PA community related to the degradation of anthropogenic pollution, and the number of genes in these pathways, are both statistically higher than those in the full set of KEGG pathways analyzed (Fishers’ Exact Test, p=0.0081 for number of pathways, p<0.001 for number of genes in the KEGG pathways in PiCRUST). The enrichment for pathways involved in degrading anthropogenic pollutants is specific for the comparison of PA vs FL: Pathways co-varying with depth are not enriched in these functions (Fisher’s Exact Test, p=0.702), and while there may be some seasonal signal to the enrichment of these pathways, it is not significant (Fisher’s Exact Test p=0.054).

Only two pathways were enriched in the heterotrophic FL bacteria compared to the PA ones, consistent with their typically smaller genomes, one for the degradation of polycyclic aromatic hydrocarbons (PAHs) and the other for the degradation of inositol phosphates. PAHs may be produced naturally but the anthropogenic input of PAHs is far larger than that from natural sources, and thus we hypothesize that the predicted enrichment of pathways to degrade PAHs in heterotrophic FL bacteria is due to their presence as a result of anthropogenic pollution (Ghosal et al. 2016). It is known that PAH’s are present in atmospheric aerosol particles deposited in the EMS particularly from air masses that have passed over polluted areas of southern Europe (Iakovides et al. 2019). Inositol phosphates are a group of ubiquitous signaling molecules in animals, a major form of P storage in plant seeds, and abundant forms of P in some freshwater phytoplankton (Turner et al. 2002). They are found widely in the environment, yet are relatively under-studied and represent a major gap in our understanding of the global P cycle (Turner et al. 2002).

## Summary and outlook

Understanding the myriad factors that determine the structure and function of marine microbial communities requires two distinct view points, operating at different scales. At the one end, oceanographic processes, driven by planetary phenomena such as the changing of the seasons, determine the physical structure of the water column (e.g. temperature, stratification). These, in turn, affect the chemical conditions such as the supply and availability of nutrients. At the other end, the physiological traits that dictate where an organism can live operate at scales of nanometers or microns. These include, for example, the affinity for nutrients, the temperature optimum of enzymes or the ability of the organism to swim, perform chemotaxis and colonize particles. While clearly simplistic, these ideas provide a framework with which to interpret and understand the observed patterns in community structure in light of physical and biological drivers (Azam and Malfatti 2007, Karl 2007, Giovannoni and Vergin 2012, Seymour et al. 2017). In this study, we aimed to bridge these two viewpoints, describing how the microbial communities in the EMS change over different scales: between PA and FL communities, over depth and across seasons. Our results suggest that, in the EMS, depth and association with particles are stronger drivers of bacterial community structure when compared to seasonality. Moreover, the PA community exhibited fundamentally different dynamics from the FL one, being less diverse and exhibiting “heterotroph blooms” where different bacterial clades dominate at specific times and depths. We note that our dataset does not have the temporal resolution to fully resolve seasonal changes (e.g. (Ward et al. 2017)), in particular as we did not sample during the maximum winter bloom.

In this study, we were able to explain less than 30% of the variability in the FL and PA community structure using “a-biotic” environmental conditions (the depth and season matrices). This is consistent with most other studies of bacterioplankton in marine environments (e.g. (Louca et al. 2016), but see (Thompson et al. 2016)). This leads us to propose that organismal interactions may have a strong impact on microbial population structure. In support of this hypothesis, 20% of the variability in the FL heterotrophic population structure could be explained only when both the depth and phytoplankton community structure matrices were included. The significant factors included pigments that are associated with prymnesiophytes and *Prochlorococcus*. Previous laboratory studies have shown that both organisms interact with co-occurring heterotrophic bacteria, and in turn are affected by them in both positive and negative ways (e.g. (Sher et al. 2011, Segev et al. 2016, Ma et al. 2017, Barak-Gavish et al. 2018)). The observation that the presence of these phytoplankton taxa correlates mostly with heterotrophic FL population structure suggest that, in the EMS, the main interactions occur through the production and utilization of dissolved organic matter rather than being contact-mediated interactions, e.g. in the phycosphere.

Aiming to raise testable hypotheses as to additional environmental factors that may impact the heterotrophic bacterial population structure in the EMS, we identified a possible enrichment in the PA community versus the FL one in pathways for the metabolism of anthopogenic pollutants, including pesticides, petroleum products and plastics. These results are based on an inference of community metagenomes, which may be biased by many factors, including unequal representation of different heterotrophic bacterial lineages in the genomic databases and genomic variability between taxa with closely related 16S sequences. We also currently do not have measurements of such pollutants across time and space from the EMS, which are critical in order to test the potential impact of such molecules. We note, however, that, possibly due to analytical reasons (costs, detection limits etc.), anthropogenic pollutants are rarely incorporated into the oceanographic “world view”. Very few oceanographic cruises or time-series observatories include measurements of such contaminants. As a result, global maps of their distributions and dynamics (e.g. similar to those for nutrients, trace metals or organic carbon) are rare or nonexistent. Nevertheless, anthropogenic pollutants can affect the physiology of marine microorganisms, providing nourishment to some groups of organisms (Romera-Castillo et al. 2018) but potentially poisoning others, including picocyanobacteria which are at the base of oligotrophic food webs (Echeveste et al. 2010, Tetu et al. 2019). Our results suggest the need for better understanding of anthropogenic pollutants as potential drivers of microbial communities in the oceans.

Finally, to what extent can the results and hypotheses from this study be generalized from the EMS to other oligotrophic oceans? The EMS is unique among oligotrophic marine environments in being strongly depleted in P, with N:P ratios of inorganic nutrients, dissolved and particulate organic matter >>16:1, compared to ^~^16 in most other marine environments (Krom et al. 2005). This might explain the enrichment in the heterotrophic FL communities in the pathway for utilization of inositol phosphates, which are relatively abundant sources of organic P yet is not well characterized in the oceans (Turner et al. 2002). Additionally, the proximity of the EMS to land might increase the input fluxes of anthropogenic pollution, although such pollution is prevalent also in the most remote areas of the ocean (Rios et al. 2010, Chen et al. 2018). Given the relative accessibility of the EMS and its importance as a sea that provides ecosystem services to millions of people, we anticipate that it will become a useful model system to study the processes affecting oligotrophic microbial communities, and how these processes change in a changing world.

## Brief Experimental Procedures

### Sampling and analysis of nutrients, cell numbers and photosynthetic pigments

Six one-day cruises were performed over a two-year period to station n-1200 (32 27.36 N, 034 22.47 E) onboard the R/V Mediterranean Explorer. Two cruises were in fall, (December 1^st^, 2014; November 18^th^, 2015), two in spring (March 24^th^, 2015; March 30^th^, 2016) and two in summer (July 14^th^, 2015; July 25^th^, 2016). Samples were collected using 8L Niskin bottles mounted on a rosette with a SeaBird CTD profiler (SBE 19plus V2). Water was collected from 5-6 depths, corresponding to surface waters (10m), one-half of the observed DCM, the DCM, 200m and 500m (during the November 2015 cruise, when there was a shallow chlorophyll maximum, samples were collected at 50m and 100m). In several cruises an additional sample was collected from below the DCM and above 200m (termed the “Twilight zone”, or TZ, Supporting Information Excel Table). Due to the amount of water needed for each analysis a separate rosette cast was performed for each depth, resulting in samples being collected up to ^~^4 hours apart. The mixed layer depth was defined as the depth where potential density differed from surface values by >0.125 kg m^−3^ (Malmstrom et al. 2010). We note that, during the spring cruises, there was no clear pycnocline and therefore the actual mixing likely extended significantly below the calculated MLD, but the estimates of the effect of seasonality on community structure are robust to differences in the MLD. Photosynthetic pigments were analyzed using a method adapted for UPLC from the LOV method (Hooker et al. 2005). More information on the pigment analysis, as well as details of the nutrient and flow cytometry analyses, can be found in the Supporting Information methods section.

### DNA sequencing and analysis

5-11.5L of seawater were filtered using a peristaltic pump onto three filters maintained in-line: 47mm 11 and 5 μm nylon filters and 0.22 μm sterivex filters (Millipore). Storage buffer (40 mM EDTA, 50 mM Tris pH 8.3, 0.75 M sucrose) was added to the samples which were frozen on-board on dry ice and maintained at −80°C until analysis. DNA was extracted using a semi-automated protocol that included manual mechanical and chemical cell lysis followed by automated nucleic acid extraction with a QIAcube system (Haber et al. 2020). PCR amplification was performed using the 16S primer set 515F-Y and 926R that targets the variable V4-5 region and is modified to amplify common oligotrophic bacterial lineages such as SAR11 (Parada et al. 2016). Libraries were sequenced using a MiSeq instrument, paired-end sequencing reads were merged, denoised, pre-processed, and assigned to taxonomic identifiers using Dada2 (version 1.1.6) (Callahan et al. 2016). Exact Sequence Variants (ESVs) were assigned taxonomy using the “classify.seqs” command in MOTHUR, the SILVA database (version 128) and an 80% identity cutoff (Schloss et al. 2009). For Variation Partitioning Analysis (VPA), we defined three matrices of conditions corresponding to those associated with seasonality (cruise number, season, sea surface temperature and mixed layer depth), depth (depth, temperature, NOx concentration and salinity) and phytoplankton community structure (the ratios of divinyll-chlorophyll a, chlorophyll b, 19’-hex fucoxanthin, fucoxanthin, peridinin and diadinoxanthin to total chlorophyll a). VPA was performed using “VarPart”, followed by conditional Canonical Correspondence Analysis (CCA) using the “cca” and “anova.cca” commands (all in the Vegan package). Metagenome inference from denoised sequences were performed using PICRUSt (Langille et al. 2013), as described previously (Goldford et al. 2018). Communities were normalized using the normalize_otus.py function in PICRUSt, and the metagenomes were estimated using the estimate_metagenome.py routine. The weighted NSTI values ranged between 0.07-0.25 (mean 0.16) for the PA heterotrophic bacteria and 0.09-0.20 (average 0.15) for the FL heterotrophs, within the range of other less-studied environments such as soil and mammalian metagenomes (Langille et al. 2013), suggesting that the results are useful to raise testable hypotheses but should be interpreted with caution. Due to the relatively large sample sizes (e.g. when comparing sample dissimilarities), Welch’s t-tests were used in Microsoft Excel to compare means, as recommended by (Fagerland 2012). More detailed information on the DNA extraction, sequencing and quality control can be found in the Supporting Information methods section.

### Data availability

An excel table with the full environmental dataset, the ESV tables with and without cyanobacteria, and the dynamics of ESVs belonging to specific clades, are presented in the Supporting Excel File. The oceanographic data were deposited in the BCO-DMO under project acronym HADFBA. The sequencing reads were deposited in the NCBI SRA database under project number PRJNA548664. These data include also a transect from the coast to station n-1200, described elsewhere, (Haber et al. 2020).

## Acknowledgments

We thank the captain and crew of the R/V Mediterranean Explorer for their invaluable assistance during the research cruises. We also thank the scientific team on all cruises for their assistance, Dr. Tanya Rivlin (Interuniversity Institute for marine sciences, Eilat) for the nutrient analyses and Dr. Stephan Green (DNA Services Facility at the University of Illinois at Chicago) for the amplicon sequencing. This study was funded by the Israel Science Foundation grant (ISF #1243/16, to LS), the Human Frontiers Science Program (grant number grant RGP0020/2016, to D. Segrè and D. Sher) and by the National Science Foundation - United States-Israel Binational Science Foundation Program in Oceanography (grant number 1635070/2016532 to D. Segrè and D. Sher). The seasonal cruises were supported by funding from the Leon H. Charney School of Marine Sciences (Haifa University, Israel) and subsidized by the EcoOcean Foundation. MH was supported by an Inter-Institutional post-doctoral fellowship from the Haifa University and a Helmsley Trust fellowship.

