## Supporting Information for "Particle-associated and free-living bacterial communities in an oligotrophic sea are affected by different environmental and anthropogenic factors"

**Supporting methods**

**Nutrient and flow cytometry analysis:** Seawater for nutrient analysis was collected in 15 ml Falcon tubes pre-rinsed with sample seawater. For each nutrient, duplicate non-filtered samples were frozen onboard directly after collection and kept at -20°C until analysis of silicate, nitrate+nitrite and soluble reactive phosphate content (within six weeks) at the service unit of the Interuniversity Institute for Marine Sciences in Eilat, Israel. Precision limits are 0.05 µmol/l for silica and nitrate+nitrite and 0.04 µmol/l for phosphate. For flow cytometry, triplicate 1.5 ml seawater samples were fixed with glutaraldehyde (0.125% final concentration), incubated in the dark for 10 min, stored in liquid nitrogen onboard and kept at -80 °C in the lab until analysis using a BD FACSCanto™ II Flow Cytometry Analyzer Systems (BD Biosciences) with and without staining using Sybr-Green.

**Photosynthetic pigment analysis using Ultra-high Performance Liquid Chromatography (UPLC):** Photosynthetic pigments were analyzed using a method adapted for UPLC from the LOV method (Hooker et al. 2005). Briefly, seawater (4-11L) was filtered on-board onto glass fiber filters (47mm GF/F, Whatman, nominal pore size 0.7μm) using a peristaltic pump, excess water was absorbed onto a kimwipe and the filters were flash-frozen on-board in liquid nitrogen and stored in the lab at -80°C until analysis. Pigments were extracted in 1 ml 100% methanol for 2.5 h at room temperature, clarified using syringe filters (Acrodisc CR, 13 mm, 0.2 µm PTFE membranes, Pall Life Sciences) and analyzed on an ACQUITY UPLC system (Waters) equipped with a C8 column (1.7 µm particle size, 2.1 mm internal diameter, 50 mm column length, ACQUITY UPLC BEH, Waters) and a photodiode array detector. The samples were preheated to 30°C and 10µl was injected and the column was maintained at 50°C. The mobile phase consisted a buffer A (70:30 (v/v) methanol and 0.5M ammonium acetate) and buffer B (100% methanol). The run protocol started with the stabilization of the system for 0.2 min to the initial mixture of 80:20 (buffer A:buffer B) and flow rate of 0.5 mL min^-1^. The run continued with 1.8 min linear gradient to 50:50, then linear gradient to 100% buffer B for 7 min, 3 min of 100% buffer B and 0.5 min linear gradient to the initial buffer proportion 80:20 that continued until the end of the run (14 min). Pigment identification and quantification was performed based on retention time and spectrum absorbance using standards for chlorophyll a, divinyl-chlorophyll a, chlorophyll b, chlorophyll c2, zeaxanthin, beta-carotene, diatoxanthin, fucoxanthin, peridinin (fucoxanthin was used to estimate the concentrations of 19’-hex-fucoxanthin and 19’-but-fucoxanthin). All standards except for chlA were purchased from the DHI, Denmark.

**DNA sample collection, extraction and 16S rDNA amplification:** 5-11.5L of seawater were filtered using a peristaltic pump onto three filters maintained in-line: 47mm 11 and 5 µm nylon filters and 0.22 µm sterivex filters **(Millipore). Storage buffer** (40 mM EDTA, 50 mM Tris pH 8.3, 0.75 M sucrose) **was added to the samples which were frozen on-board on dry ice and maintained at** -80°C until analysis. DNA was extracted using a semi-automated protocol that included manual mechanical and chemical cell lysis followed by automated nucleic acid extraction with a QIAcube system (ref* Markus BioRxiv), performed at the BioRap unit, Faculty of Medicine, Technion). The first stage PCR was performed using the 16S primer set 515F-Y and 926R that targets the variable V4-5 region and is modified to amplify common oligotrophic bacterial lineages such as SAR11 (Parada et al. 2016). The primers were modified to include the CS1 and CS2 “common sequences”, enabling a second PCR amplification to prepare libraries for Illumina sequencing (2x250 paired-end reads) using an Illumina MiSeq sequencer. Library preparation and pooling were performed at the DNA Services (DNAS) facility, Research Resources Center (RRC), University of Illinois at Chicago (UIC). MiSeq sequencing was performed at the W.M. Keck Center for Comparative and Functional Genomics at the University of Illinois at Urbana-Champaign (UIUC). Negative controls from the first PCR had ~25-50-fold fewer sequencing reads compared to the samples, suggesting cross contamination was negligible. The Bray-Curtiss Dissimilarity between replicate samples (from the same cast) was significantly lower than the dissimilarity between samples by season, depth or size-fraction (t-test, p<0.001 for all comparisons), suggesting that the observed community patterns are robust to within-sample variability.

**Sequence processing and statistical analyses:** Paired-end sequencing reads were merged using Flash (version 1.2.11) with the following parameters: max-overlap: 95; min-overlap:85, with all other parameters set to default (Magoč and Salzberg 2011). Merged reads were denoised, pre-processed, and assigned to taxonomic identifiers using Dada2 (version 1.1.6) (Callahan et al. 2016). First, reads were trimmed to 400 nucleotides, while all other parameters were set to default values. Bimeric PCR products were then removed using the “tableMethod” parameter set to “consensus.” Exact Sequence Variants (ESVs) were assigned taxonomy using the “classify.seqs” command in MOTHUR, the SILVA database (version 128) and an 80% identity cutoff (Schloss et al. 2009). Eukaryotic, archaeal and uncharacterized sequences (no assigned taxonomy) were removed, and subsampling was performed to 10,000 reads using the “rrarefy” command in the R Vegan package. For analyses of heterotrophic sequences only, cyanobacterial sequences were removed and the resulting datasets were subsampled to 9,400 reads. Non-Metric Multidimensional Scaling (NMDS) was performed using metaMDS in vegan on log10 transformed data (Bray-Curtis dissimilarity, k=3, try=100, maxiters=100). For Variation Partitioning Analysis (VPA), we defined three matrices of conditions corresponding to those associated with seasonality (cruise number, season, sea surface temperature and mixed layer depth), depth (depth, temperature, NOx concentration and salinity) and phytoplankton community structure (the ratios of divinyll-chlorophyll a, chlorophyll b, 19’-hex fucoxanthin, fucoxanthin, peridinin and diadinoxanthin to total chlorophyll a). VPA was performed using “VarPart”, followed by conditional Canonical Correspondence Analysis (CCA) using the “cca” and “anova.cca” commands (all in the Vegan package). Metagenome inference from denoised sequences were performed using PICRUSt (Langille et al. 2013), as described previously (Goldford et al. 2018). Communities were normalized using the normalize_otus.py function in PICRUSt, and the metagenomes were estimated using the estimate_metagenome.py routine. The weighted NSTI values ranged between 0.07-0.25 (mean 0.16) for the PA heterotrophic bacteria and 0.09-0.20 (average 0.15) for the FL heterotrophs, within the range of other less-studied environments such as soil and mammalian metagenomes (Langille et al. 2013), suggesting that the results are useful to raise testable hypotheses but should be interpreted with caution. Due to the relatively large sample sizes (e.g. when comparing sample dissimilarities), t-tests were used to compare means, as recommended by (Fagerland 2012).

**Supporting text – depth distribution of abundant free-living and particle-associated ESVs**

The ESV-level resolution of our data allows us to describe the dynamics of closely related organisms, defined as sharing the same V4-V5 sequence in their 16S rDNA (Figure 3). We focus here on four main clades – the free-living SAR11 clade and pico-cyanobacteria (*Prochlorococcus* and *Synechococcus*), and the particle-associated *Alteromonas* and *Ralstonia*.

**SAR11 (Pelagibacterales):** ESVs identified as SAR11 were reclassified using the Silva database 132 (accessed 24.09.2019), allowing us to assign 142 of the 166 SAR11 ESVs to clades I-IV representing between 80.6 and 100% of reads assigned to SAR11 of the FL samples (average 94.7%). This reclassification allows a direct comparison with previous studies, revealing that the spatial and temporal dynamics of the identified clades follow largely known dynamics for these clades from the Bermuda Atlantic Time Series station (BATS, (Vergin et al. 2013)) and from surface waters at an offshore northwest Mediterranean Sea station (Salter et al. 2015).

SAR11 clade I was the most abundant SAR11 clade at all times and depths contributing between 45.9 and 85.2% of SAR11 reads. Of its 82 ESVs, 59 could be further assigned to subclades Ia (16 ESVs), Ib (32 ESVs) and an uncultured clade (11 ESVs), which based on Blast analysis corresponded to subclade Ic . These subclades showed different spatial and temporal patterns.

Clade Ia was most abundant at 10m and absent from 500m in all cruises. It always contributed most SAR11 reads of all identified groups at 10m and 1/2DCM with its maximum abundance in summer at 10m. This dominance in surface waters throughout the year was also reported from an offshore station in the NW Mediterranean Sea (Salter et al. 2015), and contrast the dynamics at BATS, where clade Ib replaces Ia as the most common SAR11 group at the surface during mixing (Vergin et al. 2013). As at BATS, SAR11 clade Ib was, for the most, present at all depths throughout the year. In our study, clade Ib was more abundant than clade Ia at TZ/200m in fall. These dynamics mirror those found at BATS (Vergin et al. 2013) and at the surface also those at the NW Mediterranean (Salter et al. 2015). Clade Ic has been described as a deep ocean SAR11 bathytype discovered at BATS (Vergin et al. 2013) and ALOHA (Cameron Thrash et al. 2014). In our dataset, it was almost exclusively present at 500m where it represented the most abundant of the SAR11 subclade I.

Clade II was present at all depths year round and was the second most common SAR11 clade. It usually had its maximal abundance at 500 m. In fall, it had a second maximum at 10m and was lower at the intermediate depths. Clade II has been shown to be divided into two subclades, IIa and IIb, with different ecotypes (Vergin et al. 2013). Clade IIa blooms in spring at the surface at BATS, while clade IIb, is present at deeper depth. This difference could explain the distribution observed for clade II at station n-1200.

Clade III was only detected in two samples (10m, in fall 2015 and in summer 2016). Both times making up about 1% of the FL community. At BATS, clade IIIa has also been reported at low abundance, present in autumn surface waters at the onset of mixing (Vergin et al. 2013). SAR11 clade IV was mostly found between 10 and the DCM, while it was absent or extremely rare at 500 m.

***Prochlorococcus* and *Synechococcus*:** An analysis of the most abundant picocyanobacterial ESVs revealed dynamics consistent with seasonal dynamics of these ecotypes at the Bermuda Atlantic Time Series (BATS) station (Malmstrom et al. 2010). During stratified periods, two *Synechococcus* ESVs were the most abundant ESVs in the surface waters, consistent with the flow cytometry data showing a higher abundance of *Synechococcus* cells compared to *Prochlorococcus* at the surface water in most stratified seasons (Supplementary Figure S3). The *Synechococcus* ESVs were both most closely related to strain WH8109, and thus belong to clade II, the most abundant open ocean Synechococcus clade (Sohm et al. 2016). The *Synechococcus* ESVs were replaced at intermediate depths (1/2 DCM) with two high-light *Prochlorococcus* ESVs (closely related to strain MED4) as the dominant cyanobacteria ESVs (Figure 3C). Low-light adapted *Prochlorococcus* (related to strains NATL2A or PAC, both LLI, (Biller et al. 2014)) replaced the high-light ESVs in the DCM, with very few picocyanobacterial reads observed at 200m or 500m. During periods of mixing (the two spring cruises) the *Synechococcus* and high-light *Prochlorococcus* ESVs were more-or-less homogenously distributed in the mixed layer. Additional picocyanobacterial ESVs were observed, at relatively low abundances.

**Other FL bacterial lineages:** Additional abundant ESVs belonging to the NS5 marine group (Bacteroidetes), the AEGEAN-169 clade (Rhodospirillales, α-proteobacteria) and *Candidatus* Actinomarina (OM1 clade, Actinobacteria) were common in the surface waters (10m) in each cruise with little seasonal variation. In the deeper (below 200m) water they were replaced by an abundant ESV belonging to the SAR324 clade (δ-proteobacteria).

***Alteromonas***: In about 20% of the PA samples (primarily from the summer and spring of 2015) four ESVs identified as *Alteromonas macleodii* comprised up to 65% of the sequences (Figure 3C). *Alteromonas* are copiotrophic bacteria with extremely flexible metabolism (Pedler et al. 2014, López-Pérez and Rodriguez-Valera 2016), which often increase in numbers or activity in bottle incubations (e.g. in response to DOM amendments, but often also in controls (McCarren et al. 2010, Shi et al. 2012)). While they are occasionally observed in amplicon libraries to comprise a significant portion of the reads (Acinas et al. 1999), they have been observed only rarely by FISH (Eilers et al. 2000), leading to their ecological importance being questioned. However, RNA sequences from *Alteromonas* are common in deep-sea transcriptomes, suggesting they may be have a significant role in recycling of organic carbon and nitrogen, e.g. from amino acids (Baker et al. 2013). Our results further support the notion that *Alteromonas macleodii* can be a dominant organism in particle-associated communities under oligotrophic conditions, primarily during spring and summer. Combined with observations that *Alteromonas* may be relatively active in natural samples (e.g. higher relative abundance of RNA transcripts *vs.* DNA sequences, even in samples of FL bacteria (Shi et al. 2010, Baker et al. 2013)), these results suggest that this clade may dominate both in numbers and in activity at specific times and locations.

Interestingly, ESVs belonging to another species of *Alteromonas*, *A. mediterranea*, were also observed but at consistently lower abundances*. Alteromonas mediterranea* was previously designated as a “deep ecotype” of *A. macleodii* (López-Pérez and Rodriguez-Valera 2016), yet in our dataset there is no clear depth partitioning between these organisms (Supplementary excel file). Whether or not *A. macleodii* is always more common, and whether there is a clear niche partitioning between these organisms in the EMS, remains to be determined.

***Ralstonia***: While *Alteromonas* were dominant primarily in spring and summer, *Ralstonia* (β-proteobacteria) were dominant in PA samples taken during fall 2015, primarily in deeper samples where two ESVs comprised up to ~33% of the population. The sequences identified were most similar to *Ralstonia picketii*, which can live under oligotrophic conditions (Ryan and Adley 2013). In the context of marine environments, *Ralstonia picketii* have been recorded primarily in association with fish or cnidarians (Matyar 2007, Schuett et al. 2007, Garren et al. 2008). While *Ralstonia* sequences are often identified as contaminants in 16S amplicon sequencing projects (Salter et al. 2014), they were not common in our negative controls, and the DNA concentration in the samples from which these bacteria were identified were not especially low. Thus, the presence of these organisms in the PA populations from the fall 2015 and, to a lesser extent, in FL populations from the same time, suggests they were relatively common members of the microbial community at this time.

**Additional PA bacterial lineages**: Additional organisms that were dominant in specific PA communities were *Sphingobium* (~18% of the community at 130m during fall 2014), *Oleibacter* (~40% during spring 2016 at 50m and up to ~30% during the summers of both 2015 and 2016 at 10m) and *Acinetobacter* (~18% during fall 2014 at the DCM). Strains from all of these genera are known for their ability to degrade petroleum products or PAHs (Teramoto et al. 2011, Ghosal et al. 2016).

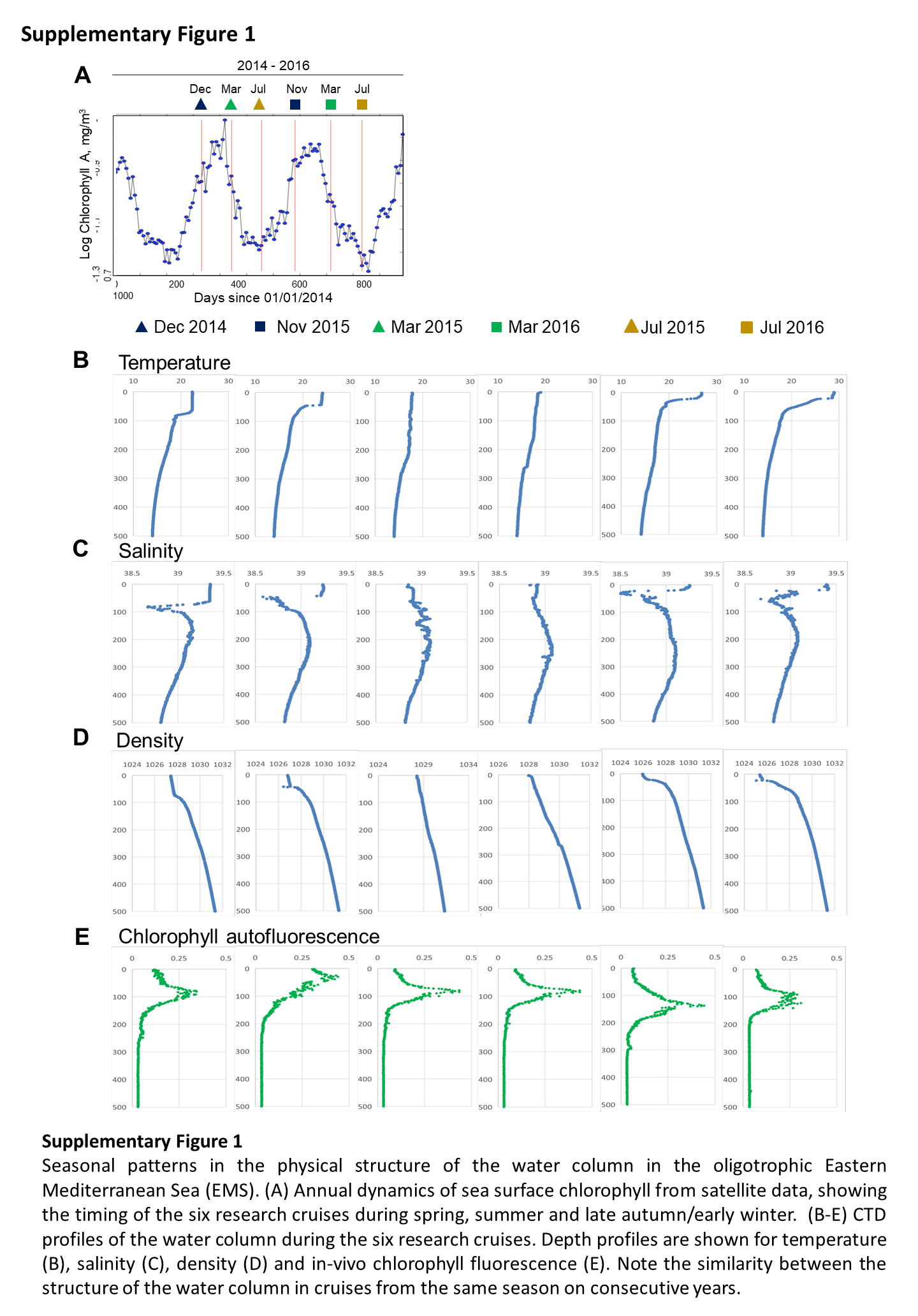

**Supporting Information Figure S1:** Seasonal patterns in the physical structure of the water column in the oligotrophic Eastern Mediterranean Sea (EMS). (A) Annual dynamics of sea surface chlorophyll from satellite data, showing the timing of the six research cruises during fall, spring and summer. (B-E) CTD profiles of the water column during the six research cruises. Depth profiles are shown for temperature (B), salinity (C), density (D) and *in vivo* chlorophyll fluorescence (E). Note the similarity between the structure of the water column in cruises from the same season on consecutive years.

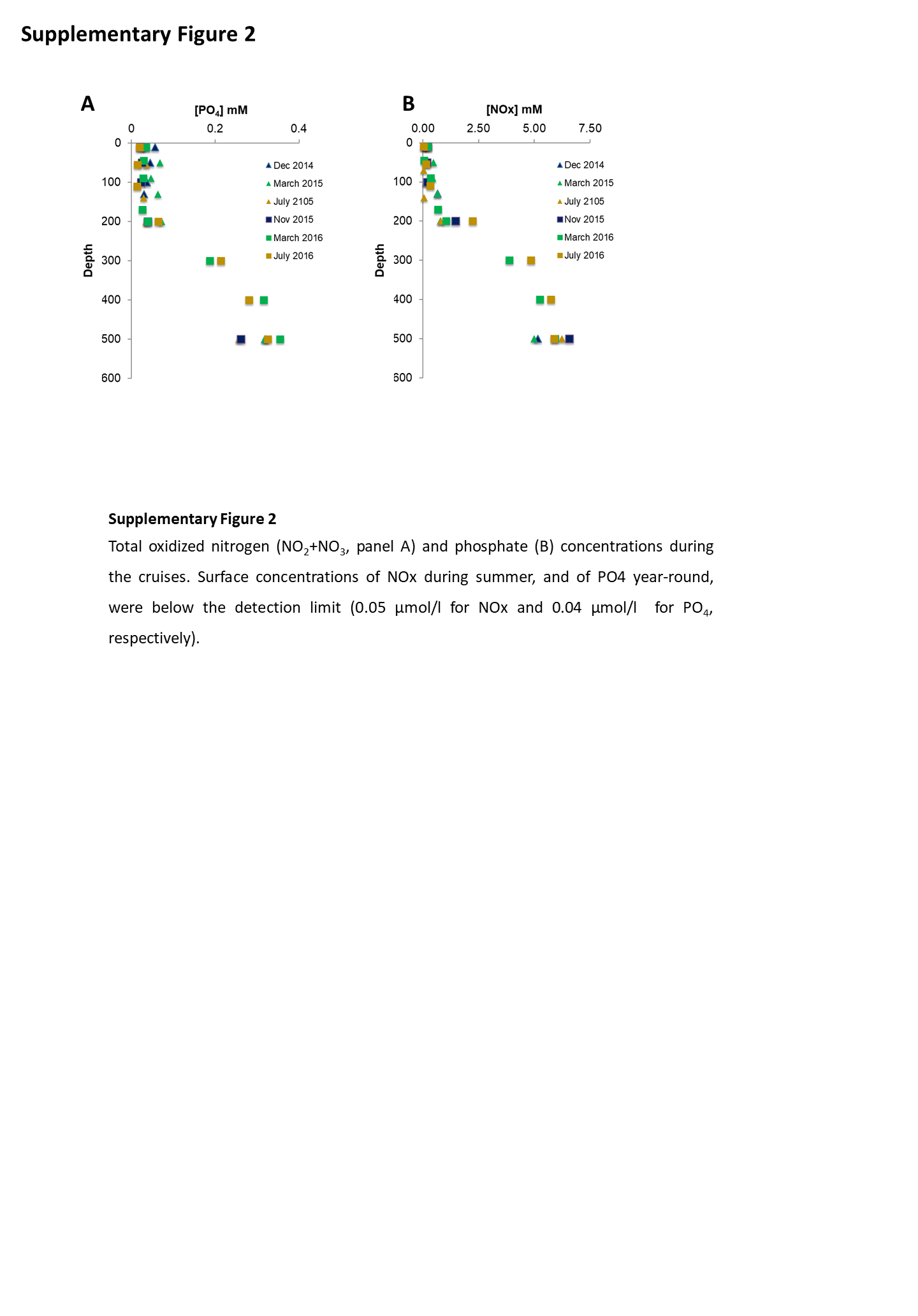

**Supporting Information Fig. S2:** Total oxidized nitrogen (NO_2_+NO_3_, panel A) and phosphate (B) concentrations during the cruises**.** Surface concentrations of NOx during summer, and of PO4 year-round, were below the detection limit (0.05 µmol/l for NOx and 0.04 µmol/l for PO_4_, respectively).

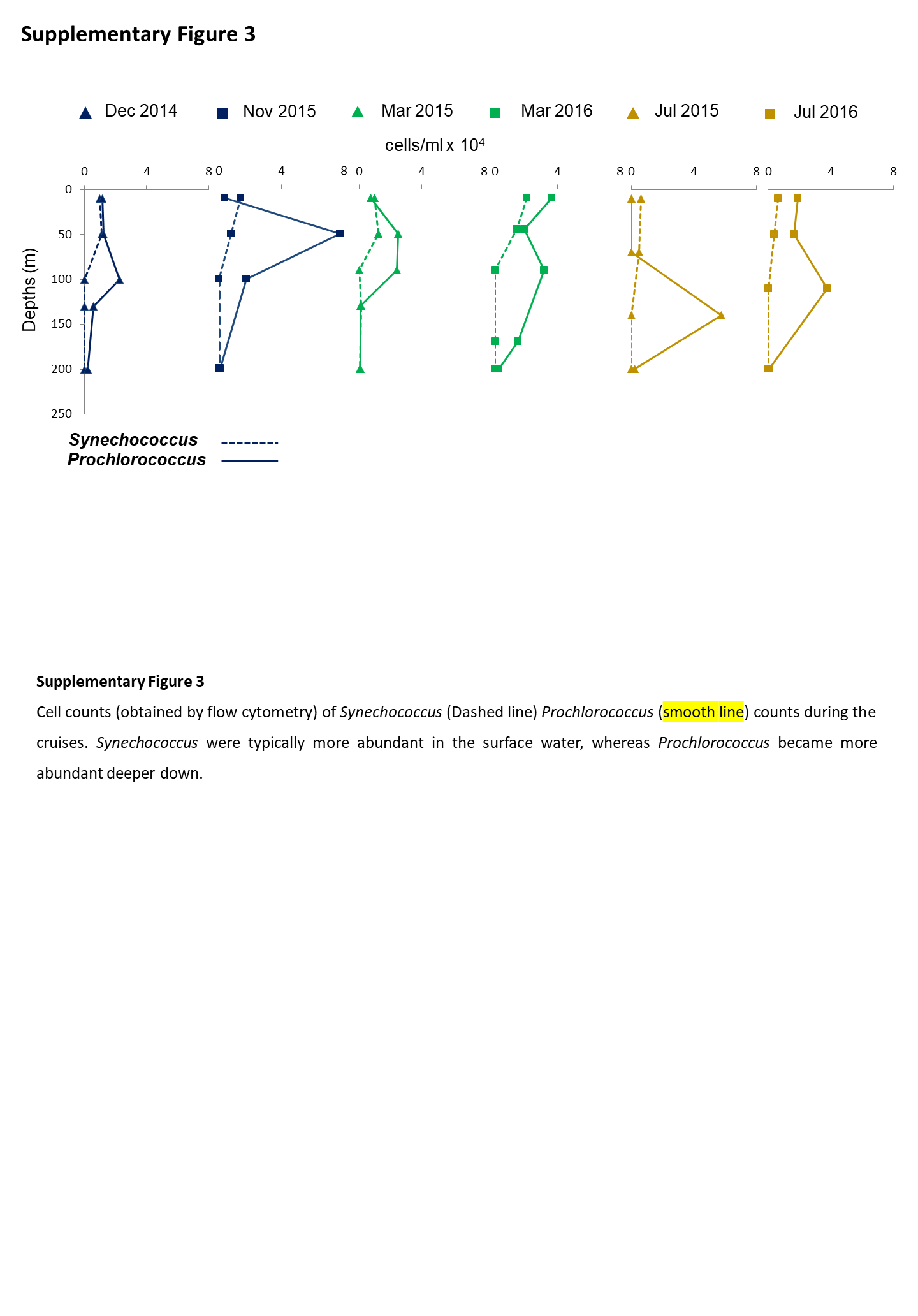

**Supporting Information Fig. S3:** Cell counts (obtained by flow cytometry) of *Synechococcus* and (dashed line) *Prochlorococcus* ( continuous line) counts during the cruises. *Synechococcus* were typically more abundant in the surface water (10 m), whereas *Prochlorococcus* became more abundant deeper down (especially at the DCM). Data points represent averages of three technical replicates. Note that the depths of the 1/2DCM and DCM change over the cruises, with the ½ DCM typically around 50m and the DCM between 90-130m.

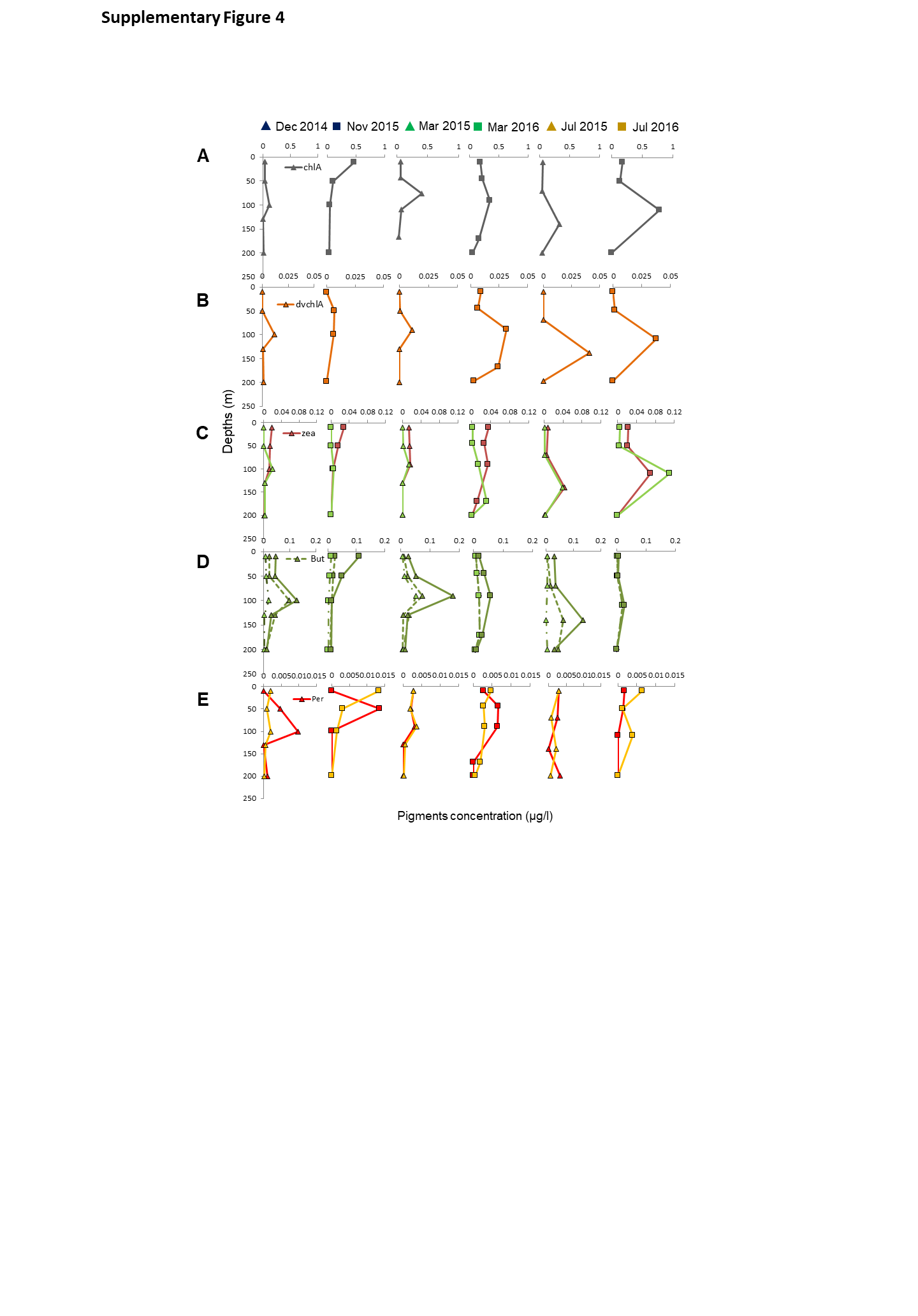

**Supporting Information Figure S4:** Photosynthetic pigment concentrations during the cruises. Chlorophyll-a (panel A) is found in all photosynthetic organisms apart from *Prochlorococcus*, which contains divinyl-chlorophyll a (dvchlA) instead (panel B) (Veldhuis and Kraay 1990, Moore et al. 1995). Zeaxanthin and chlorophyll B (zea, chlB, respectively) are found in multiple phytoplankton but are often associated primarily with cyanobacteria (Jeffrey et al. 2005). Fucoxanthin (Fuco) and Diadinoxanthin (Diadino) are associated mainly with diatoms (D-E) but appear also in other phytoplankton such as prymnesiophytes. 19’-hex-fucoxanthin is considered a marker of prymnesiophytes, whereas 19’-but-fucoxanthin (but) is also found in Pelagophytes and diatoms, and is thus less specific (Jeffrey et al. 2005). Peridinin is associated with dinoflagellates (E). Note the differences in scale between the pigments. Note that the depths of the 1/2DCM and DCM change over the cruises, with the ½ DCM typically around 50m and the DCM between 90-130m.

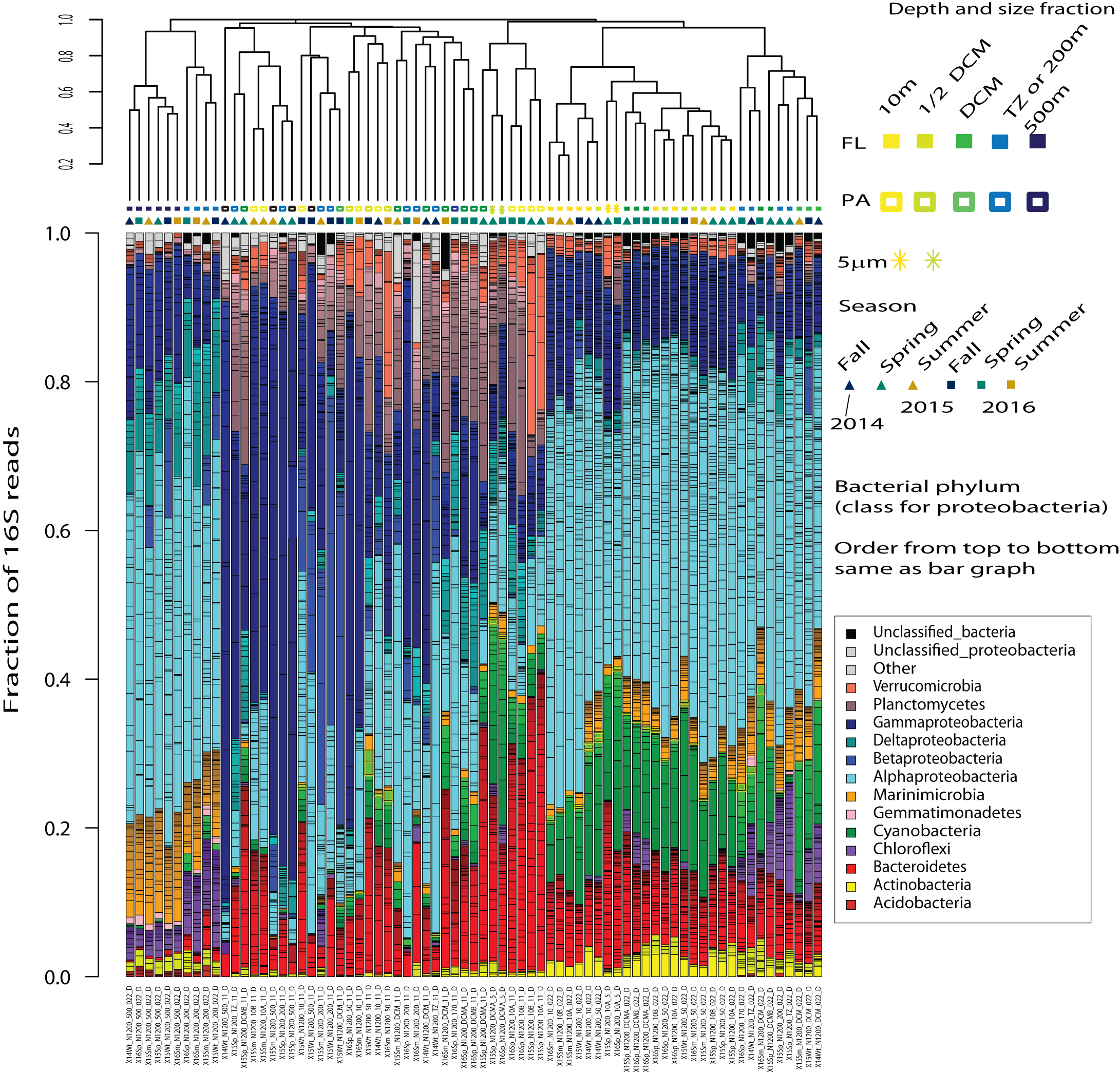

**Supporting Information Figure S5:** Bar graph of the microbial communities. Different colors represent bacterial phyla (or, in the case of Proteobacteria, different classes). Black lines within each phylum represent specific ESVs. The samples are ordered by Bray-Curtis dissimilarity. Colored shapes between the cladogram and the bar plot represent depth and size fraction (upper row) and season (lower row).

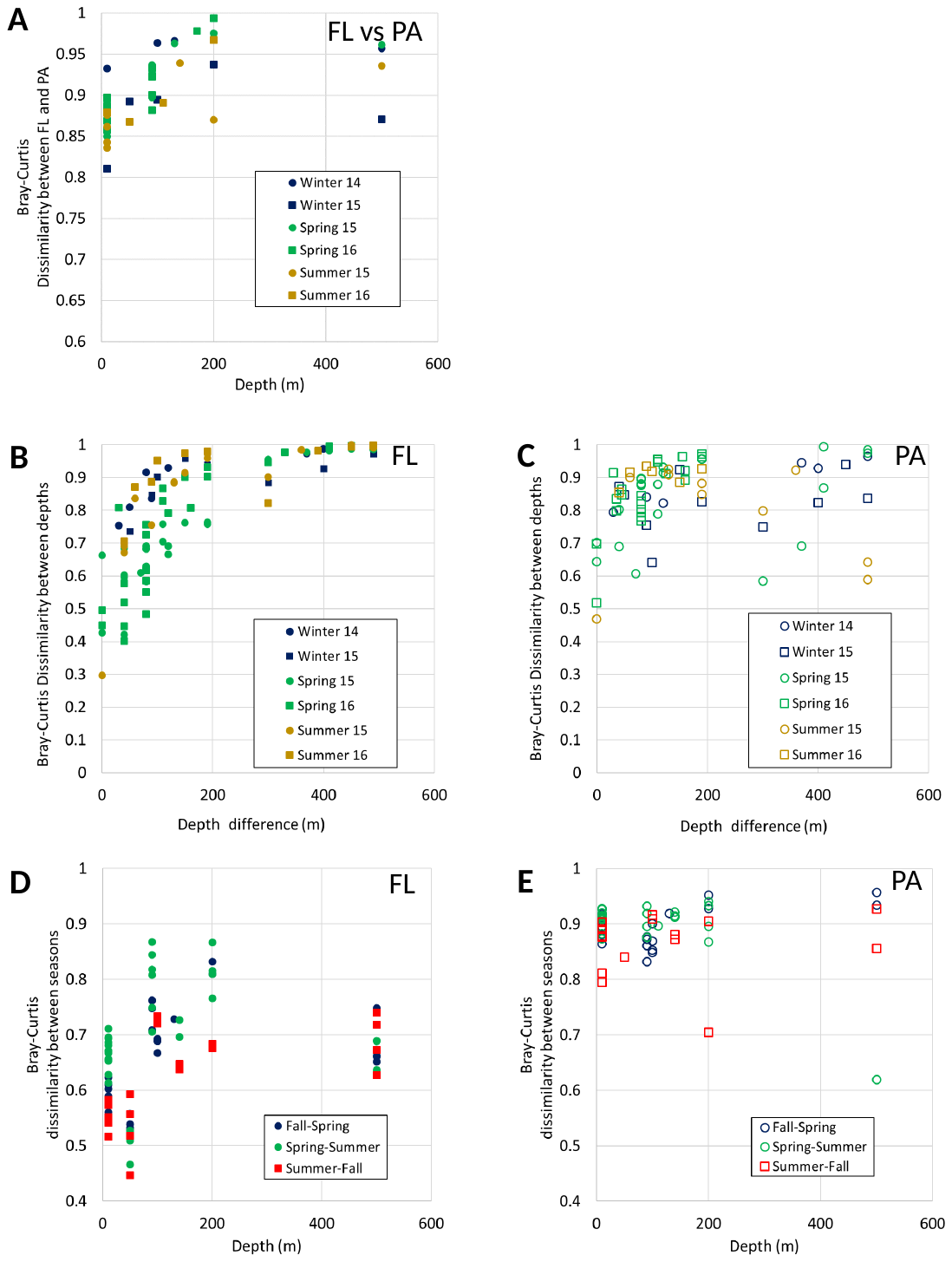

**Supporting Information Figure S6:** Distances between samples by depth, size fraction and season. In all panels, the Bray-Curtis dissimilarity between pairs of samples is plotted. The dissimilarity was calculated within groups, e.g. comparisons of depth were performed within the same cruise and size fraction, etc. (A) Dissimilarity between FL and PA samples increases with depth, with no clear seasonal trend. (B, C) Dissimilarity between communities increases with increase in sampling depth difference in FL (B) but not PA communities (C). Additionally, in FL communities, dissimilarities with depth are higher during summer and fall, when the water column is stratified, compared to spring mixed conditions (t-test, p<0.001). D, E) Seasonal differences are higher in PA compared to FL populations. Within the FL (but not PA) community, seasonal differences are lower at the surface and 50m compared to deeper samples (t-test, p<0.001). Since the depth of the DCM changed between cruises, the depth plotted is of one of the two cruises.

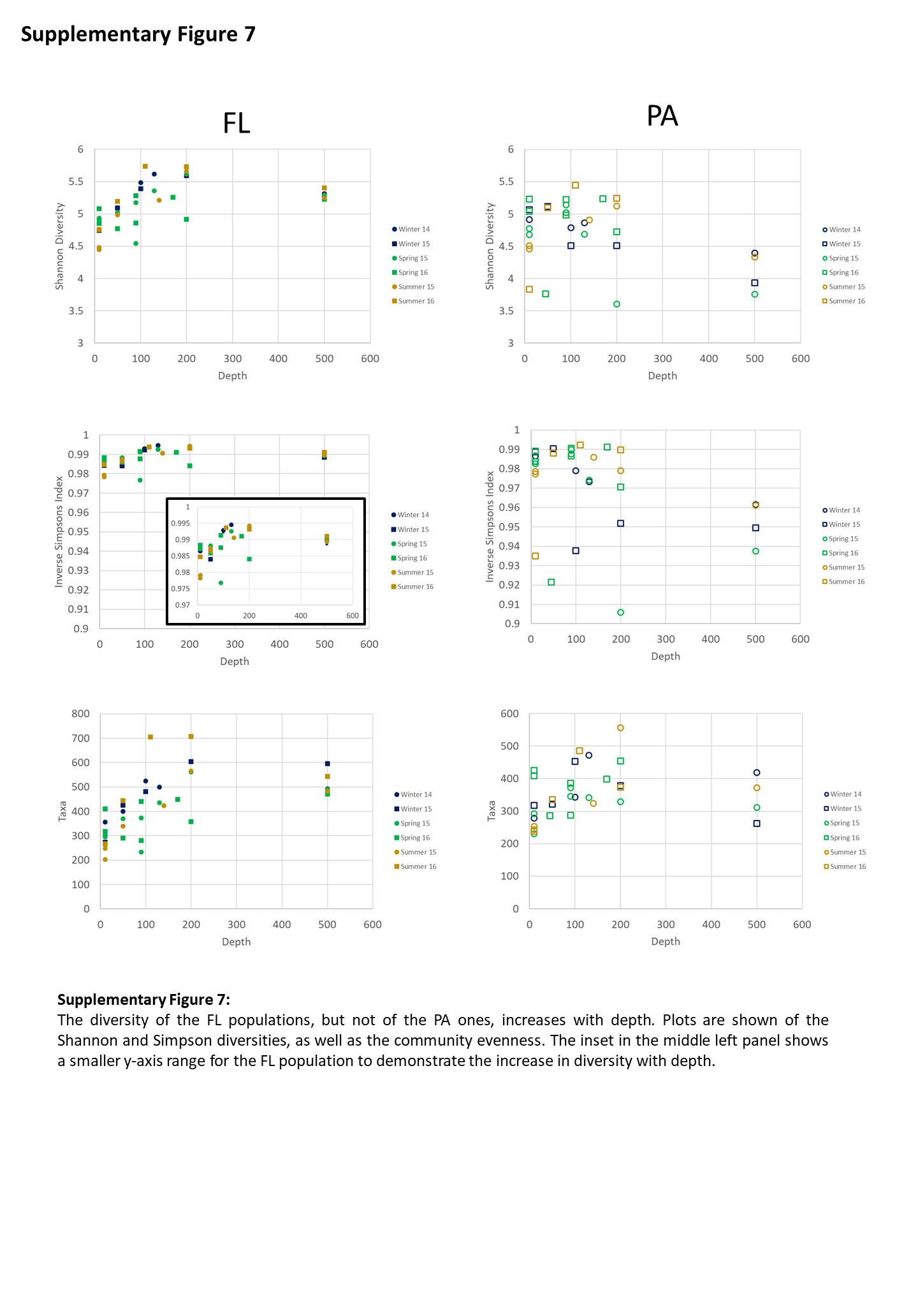

**Supporting Information Figure S7:** The diversity of the FL populations, but not of the PA ones, increases with depth. Plots are shown of the Shannon and Simpson diversities, as well as the number of observed taxa (ESVs). The inset in the middle left panel shows a smaller y-axis range for the FL population to demonstrate the increase in diversity with depth.
